# ABCG5 ABCG8-independent mechanisms fail to maintain sterol balance in mice fed a high cholesterol diet

**DOI:** 10.1101/2025.02.11.637687

**Authors:** Garrett B. Anspach, Isha Chauhan, Erika L. Savage, Brittney Poole, Victoria P. Noffsinger, Xiaoming Fu, Zeneng Wang, Ryan E. Temel, Scott R. Gordon, Robert N. Helsley, Gregory A. Graf

## Abstract

The ABCG5 ABCG8 (G5G8) sterol transporter opposes the accumulation of dietary xenosterols, but is also the primary mediator of biliary cholesterol secretion in the cholesterol elimination pathway. In humans and in mouse models of disrupted biliary cholesterol secretion, fecal neutral sterols remain constant, indicating the presence of a G5G8-independent mechanism for cholesterol excretion. Transintestinal cholesterol elimination (TICE) is thought to compensate for biliary G5G8 insufficiency. We sought to measure the compensatory increase in intestinal cholesterol secretion and provide mechanistic insight for how TICE maintains sterol balance in the absence of G5G8. Differences were not observed in fecal neutral sterols between control, acute, and chronic liver-specific G5G8 deficient mice (G5G8^LKO^). Cholesterol content did not differ at any point along the intestinal tract between genotypes. We also observed no change in the expression of apical or basolateral sterol transporting enzymes in the proximal small intestine. We then measured biliary and intestinal cholesterol secretion rates using cholesterol free and cholesterol enriched bile acid micelles as acceptors. While biliary cholesterol secretion was reduced, the intrinsic rate of intestinal cholesterol secretion did not differ between genotypes. G5G8^LKO^ and whole-body G5G8-deficient mice were challenged with a cholesterol-containing diet. While control mice upregulate fecal neutral sterol excretion, G5G8^LKO^ and G5G8^-/-^ mice fail to do so and accumulate sterol in the liver and plasma. These studies indicate that while G5G8-independent mechanisms can mediate cholesterol excretion, TICE is not upregulated in response to a loss of hepatic G5G8 and is unable to compensate for hepatic or whole-body G5G8-deficiency in response to dietary cholesterol in mice.

## Introduction

G5G8 is an obligate heterodimer expressed in the liver and intestine that promotes the secretion of sterols into bile and the intestinal lumen, respectively (1–6). Mutations in either half-transporter cause sitosterolemia (*STSL*, OMIM #210250 and #618666), a rare recessive disorder characterized by the accumulation of non-cholesterol sterols (xenosterols) in plasma and tissues (7, 8). The clinical presentation of the disease is varied and includes hemolytic disorders and thrombocytopenia, but not necessarily hypercholesterolemia (reviewed in (9)). However, multiple GWAS studies have linked rare and common variants in *ABCG5* and *ABCG8* to elevated plasma levels of total and LDL cholesterol and coronary artery disease (CAD), suggesting that while *STSL* is a recessive disorder loss of function variants or *ABCG5 ABCG8* haploinsufficiency contributes to elevated plasma cholesterol and CAD risk (10)

An estimated 70% of the body’s cholesterol is synthesized in the liver and delivered to peripheral cells in apolipoprotein B-containing lipoproteins. Non-steroidogenic peripheral cells lack the ability to metabolize excess cholesterol, thereby necessitating a pathway for its return to the liver for metabolism to primary bile acids or direct secretion into bile and elimination from the body (11). G5G8 is the principal mediator of biliary cholesterol secretion in the final step of the reverse cholesterol transport (RCT) pathway and accounts for 70-80% of biliary cholesterol secretion (2, 5, 6). However, genetic inactivation of *Abcg8* in mice does not result in a reduction in fecal sterol excretion (12). Similarly, disruptions in biliary cholesterol secretion via inactivation of bile acid secretion or bile duct ligation do not reduce, and in some cases increase fecal neutral sterols. This alternate pathway for elimination of excess cholesterol has been termed transintestinal cholesterol elimination or excretion (TICE)(13).

Stable isotope studies suggest TICE accounts for approximately 30% of whole-body cholesterol excretion in mice maintained on standard rodent diets (14). However, TICE can be increased by a number of pharmacological stimuli including statins, fibrates, and agonists of the liver-X and farnesoid X receptors (14–18). Under these conditions, TICE can account for greater than 60% of cholesterol excretion in mice. Evidence also supports a role for TICE in humans where it is estimated to account for 35% of cholesterol excretion (16, 19). However, the mediators of intestinal cholesterol uptake from the plasma compartment and G5G8-independent transport across the apical surface of intestinal enterocytes have remained elusive. The contribution of LDL, HDL, and their receptors on the basolateral surface have produced mixed results (13, 16, 20–24). While the complete absence of G5G8 has no effect on sterol balance, it was shown to be essential for pharmacologically stimulated TICE (14, 25). While the role of G5G8 in opposing dietary xenosterol accumulation is clear, the dispensability of G5G8 in cholesterol metabolism is less certain.

Either hepatic or intestinal G5G8 is sufficient to prevent the accumulation of dietary xenosterols, but the tissue-specific role of G5G8 in the maintenance of sterol balance has not been investigated (26). In the present study, we characterized the adaptive response in the intestine to hepatic G5G8 deficiency that maintains fecal neutral sterol excretion. Hepatic G5G8 was inactivated in mice using a germline liver-specific cre recombinase transgene and acutely using an adenoassociated viral vector. Following acute hepatic G5G8 deletion, the maintenance of fecal neutral sterol excretion was immediate. Intestinal cholesterol content was not reduced in the proximal small intestine, despite the reduction in biliary cholesterol secretion. In perfusion studies, the intrinsic rate of intestinal cholesterol secretion did not differ between genotypes. Finally, we challenged whole-body and liver-specific G5G8 deficient mice with a cholesterol containing diet. TICE failed to maintain fecal excretion of cholesterol with compromised biliary cholesterol secretion or whole-body G5G8-deficiency. These studies indicate that the loss of hepatic G5G8 does not result in the upregulation of TICE to maintain cholesterol excretion in mice maintained on standard rodent diets and that TICE is incapable of maintaining fecal cholesterol excretion in mice challenged with a high cholesterol diet.

## Materials and Methods

### Animal Care and Use

All animal procedures were approved by and performed under the supervision of the University of Kentucky Institutional Animal Care and Use Committee. The colony is maintained in a specific pathogen-free, temperature-controlled (21°C) facility with 14/10 light/dark cycle. Mice were group housed in individually ventilated cages with P.J. Murphy Coarse SaniChip bedding with free access to a cereal-based diet (2918 Teklad Irradiated Rodent Diet (18% protein, 6% fat, 44% carbohydrate kcal/wt) and water unless otherwise indicated. Experimental mice were generated using a trio mating scheme, weaned between 18-21 days of age, and enrolled in experiments between 8-12 weeks of age. Male and female littermates were included in all experiments unless otherwise indicated. All mice were monitored for appearance of coat condition, weight, and mobility on a weekly basis. Experimental mice were randomized to treatment groups and groups were deidentified during biochemical analyses. Conventional *Abcg5 Abcg8*-deficient mice (strain #004670) and mice harboring lox-p sites flanking exon 1 of *Abcg5* and exon 1 & 2 of *Abcg8* (*Abcg5 Abcg8*^fl/fl^ strain #026702) are maintained in our breeding colony and undergo routine strain refreshment against the C57Bl6/J strain (strain #000664, The Jackson Laboratory, Bar Harbor, ME) every 5-6 generations.

### Acute and Chronic G5G8 deficiency

*Abcg5 Abcg8*^fl/fl^ female mice were crossed with either the C57Bl6/J males harboring a Cre recombinase transgene driven by the mouse albumin promoter (strain #003574) to generate Control and G5G8 liver-specific G5G8 knockout mice (G5G8^LKO^). To accomplish acute inactivation of G5G8 (G5G8^LKO-A^), *Abcg5 Abcg8*^fl/fl^ mice were administered a control adenoassociated viral vector (AAV, serotype 2/8) or an AAV8 encoding Cre recombinase under the control of the thyroxine binding globulin promoter (AAV8_TBG-Cre, Penn Vector Core) at doses of 5x10^11^ gc/25g mouse via tail vein. At 8 weeks of age, mice were single housed in wire bottom cages and feces collected 3 days prior to and on days 3, 5, 7, 14, and 28 following AAV delivery. On day 28, mice were anesthetized under 3% Isoflurane (Covetrus 11695-6777-2), the common bile duct ligated, the gall bladder cannulated and diverted to a collection tube for 30 min (basal bile). Mice were euthanatized via cardiac puncture and liver and 5 equal segments (∼6cm) of small intestine were excised, flash frozen, and stored at -80°C until analysis. An independent cohort of Control and G5G8^LKO^ mice was euthanized at 12 weeks of age in the post prandial phase (“lights-off” + 6 hours) and 5 equal segments of the small intestine, cecum and colon and their contents were dissected and immediately extracted for lipid analysis.

Independent cohorts of mice comprised of Control and G5G8^LKO^ and wild-type (WT) and conventional *Abcg5 Abcg8* deficient (G5G8^-/-^) mice were sequentially fed purified diets in the absence and presence of excess cholesterol. Control and G5G8^LKO^ mice are maintained on standard rodent diet. At 8 weeks of age the mice were fed a custom plant sterol free (PS-Free) diet (# D10040301, 21 kcal% protein, 61 kcal% carbohydrate, 18 kcal% fat (lard)) for a period of 2 weeks. Mice were then fed this same diet supplemented with 0.2% cholesterol (w/w). Feces were collected for the final three days on each diet. To avoid confounding effects of bioactive phytosterols that accumulate in mice lacking *Abcg5 Abcg8*, this strain is maintained on the PS-free diet. Baseline feces were collected and the mice placed on the cholesterol supplemented PS-free diet for two weeks. The mice were then placed on standard diet for two weeks. As with the floxed strain, feces were collected for the final three days on each diet. At termination, basal bile was collected and tissues dissected.

### Simultaneous determination of biliary and intestinal cholesterol secretion

Secretion of cholesterol from the liver and intestine was determined as described previously (15, 24). Briefly, mice were anesthetized, the common bile duct ligated and bile diverted to collection tubes. Basal bile was collected for 30 minutes to deplete the endogenous bile acid pool. During that collection period, the tail vein was fitted with a tail vein catheter connected to a syringe pump. The proximal 10cm of the small intestine was fitted with in-flow and out-flow catheters, flushed with Krebs Henseleit Buffer (KHB), and connected to a peristaltic pump. Following basal bile collection, taurocholate (TC) was infused (100 nmol/min) and the proximal small intestine perfused with KHB containing bile acid micelles (10 mM TC, 2 mM PC, ± 0.32 mM cholesterol) at a rate of 14 µl/min.

### Analysis of mRNA and protein

The relative abundance of transcripts was determined by real-time PCR as previously reported (Tabl1 S1)(15, 27). RNA was isolated from ∼100 mg of tissue using RNA Stat-60 phenol chloroform (TEL TEST CS502; Fisher Scientific NC9489785) and purified using the Qiagen RNeasy Mini Kit (Qiagen 74106). 250ng of RNA was converted to cDNA using the iScript cDNA synthesis kit (BIO-RAD 170-8891), cDNA was diluted 1:40, and rtPCR performed with primer pairs directed towards the indicated transcript (Table S1) and the SYBR Green detection system (Life Technologies, 4364346). Data were normalized to threshold cycle values for GAPDH and HPRT and the mean of the indicated control group and expressed as fold-change by the ΔΔCt method. The relative abundance of Abcg5 and Abcg8 protein were determined by immunoblot analysis using validated antibodies developed in-house (28, 29).

### Analysis of hepatic, plasma, and biliary lipids

Hepatic lipids were extracted, resuspended in 1% Triton X100, and cholesterol content determined using commercial assay as previously described (27). Total plasma and biliary cholesterol were determined using a commercially available enzymatic colorimetric assay (FujiFilm Wako). Bile was diluted 10-fold and bile acids measured using a previously published stable isotope dilution LC-MS/MS methodology with slight modification (30). A C18 column (Kinetex^®^ 2.6 μm Polar, 50 × 2.1 mm, Phenomenex) was used to separate bile acids, with a binary gradient consisting of 0.1% propionic acid in water and 0.1% acetic acid in methanol.

### Fecal and intestinal cholesterol

Promptly after excision from mouse, whole tissue (including lumenal content) was collected in pre-weighed glass tubes primed with internal standard of 5-α cholestane. Tubes were reweighed with tissue, and Folch reagent (2:1 chloroform: methanol, 2 mL), and KOH (50%, 200µL) was added. The solution was incubated at 37°C overnight and vortexed periodically. Hexanes (2 mL) were added to the tube and vortexed, followed by addition of water (2 mL), vortexed, and centrifuged for 10 min at 10,000 rpm to separate aqueous and organic phases. The top organic phase was transferred to a clean vial and the aqueous phase was reextracted with hexanes (2 mL). An aliquot of total tissue extract (100uL) was dried down and N-methyl-N-(trimethylsilyl)trifluoroacetamide (MSTFA, 30uL) was added and heated at 95°C for 1 hour. Ethyl-acetate (70uL) was added and analyzed by injecting 5µL of sample onto a HP-5MS (0.250-mm inner diameter × 30 m × 0.25 μm) ultra-inert gas chromatography column (Agilent 19091S-433UI) at 250°C and installed in an Agilent Technologies 7890B gas chromatograph equipped with a Agilent Technologies 7693A autosampler using on-column injection and a Mass Selective Detector (Agilent G7081B). Cholesterol was quantified using AUC normalized to 5α-cholestane.

Feces were dried, weighed, and ground to a fine powder. ∼100mg of feces was added to glass tubes primed with 25 µg 5-alpha-cholestane and FNS were extracted in 2 mL of 2:1 chloroform: methanol, and 200µL of 50% KOH was added. The solution was incubated at 37°C overnight and vortexed periodically. Two mL of hexanes were added to the tube and vortexed. Two mL of water were added, sample vortexed, and centrifuged for 10 min at 10,000 rpm for phase separation. The top organic phase was transferred to GC vials, dried resuspended and analyzed by GC-MS.

### Data Analysis

Unless otherwise indicated, data are mean ± SEM. For most measures, individual values are shown in addition to summary statistics. Data were analyzed by two-way ANOVA using genotype and sex as factors with Sidak’s post-hoc tests to determine effect of genotype within sex or intestinal segment as indicated. All analyses included tests for normality. Where main effects of sex or a sex by genotype interaction were detected in the model, data were analyzed irrespective of sex using a student’s t-test. Males are shown in filled symbols and females denoted by open symbols. For intestinal and cholesterol secretion rates, cumulative cholesterol secretion was analyzed by simple linear regression within each sex and the effect of genotype determined by difference in slope.

## Results

Multiple animal models of impaired biliary cholesterol secretion maintain fecal neutral sterol output, supporting a role for enhanced TICE as an adaptive response. We compared fecal neutral sterol excretion between control, chronic and acute inactivation of hepatic G5G8 to determine the rapidity of this adaptive response. Hepatic G5G8 was deleted in *Abcg5 Abcg8*^fl/fl^ mice using an albumin-Cre transgene or acutely using AAV8_TBG-Cre and mice maintained on standard rodent diet. Hepatic, but not intestinal mRNAs for G5 and G8 were reduced in the liver of both models compared to control mice (Figure 1A). Similarly, hepatic G5 and G8 protein were reduced in both models (Figure 1B). Differences due to hepatic genotype were not observed for physiological measures such as body weight, liver weight, and basal bile flow (Table 1). As expected, biliary cholesterol concentrations and cholesterol secretion rates in basal bile were substantially reduced in mice lacking hepatic G5G8 in both males and females (Figure 1C).

**Figure 1.**
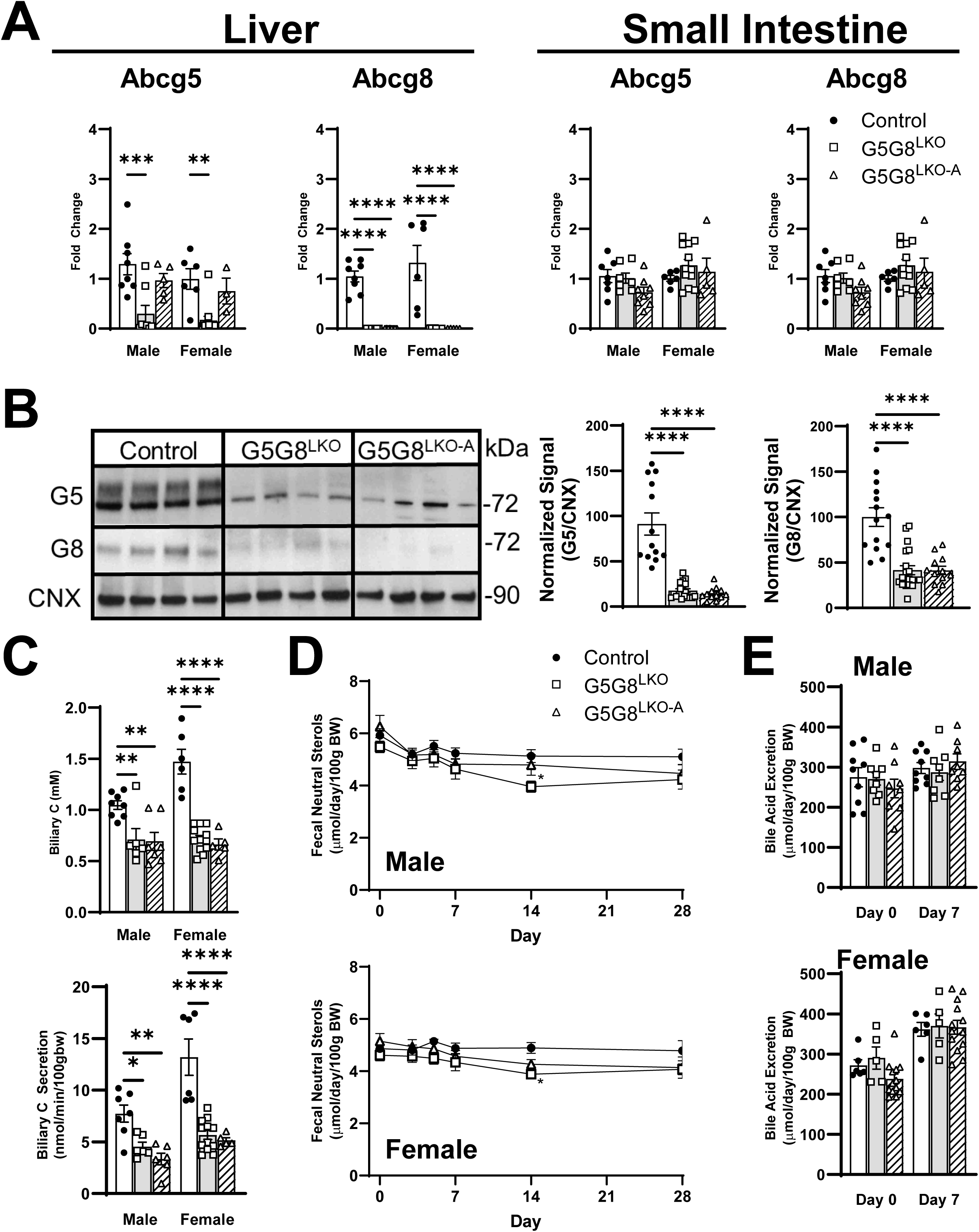
The impact of chronic or acute deletion of Abcg5 Abcg8 on biliary cholesterol secretion and fecal sterol excretion. Mice harboring floxed *Abcg5 Abcg8* were bred to the Alb-Cre strain (G5G8^LKO^) and maintained on standard diet to 8-10 weeks of age. Alb-Cre negative mice were delivered control AAV (Control) or AAV_TBG-Cre (G5G8^LKO-A^) to accomplish acute inactivation of hepatic G5G8. All mice were euthanized 28 days following AAV delivery. A) Relative abundance of Abcg5 and Abcg8 mRNA in liver and proximal small intestine. B) Detection of Abcg5 and Abcg8 protein in liver by immunoblot analysis. C) Biliary cholesterol concentrations in gallbladder bile and biliary cholesterol secretion rate in basal bile at termination of the experiment. D, E) Fecal neutral and acidic sterols prior to (Day 0) and up to 28 days following AAV delivery. Data are mean ± S.E.M (n=6-12) in males and females and analyzed by two-way ANOVA using sex and genotype as factors. Effects of genotype within sex were determined using a Sidak’s multiple comparison tests. Sequential measures of fecal neutral sterols (D) were conducted within sex using a two-way repeated measures ANOVA using genotype and time as factors. A Dunnet’s post-hoc test was used to determine differences from Day 0. *<0.05 **P<0.01 ***P<0.001 ****P<0.0001.

**Table 1.**
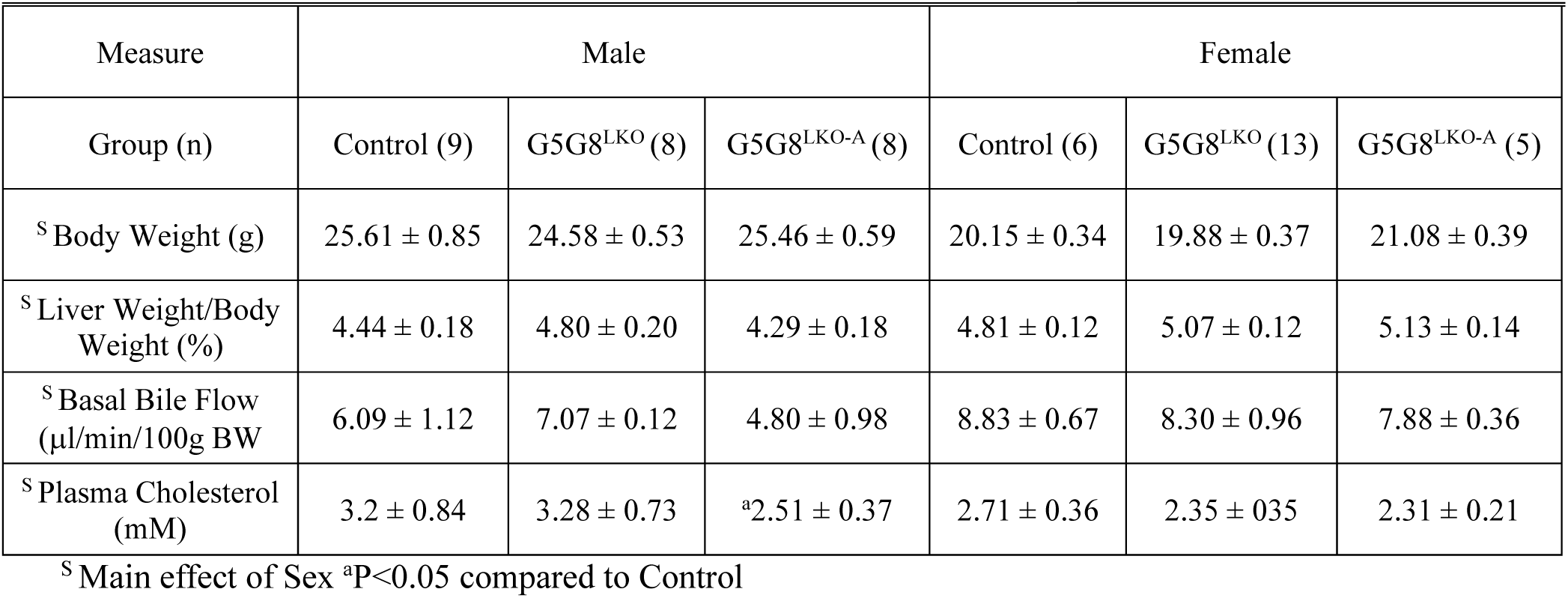
Physiological Measures in Control and liver-specific G5G8-deficient mice.

Baseline fecal neutral sterols did not differ between groups (Figure 1D). Nor did they differ for up to 28 days between controls and mice with either chronic or acute inactivation of G5G8. The apparent downward trend in FNS over the 28-day period, particularly in males, is largely due to weight gain during this period as the data are normalized to body weight. We also did not observe differences in fecal bile acids between groups at the 28-day time-point (Figure 1E). These data indicate that the TICE can adapt to the loss in hepatic G5G8 within a timescale not discernable under these experimental conditions. In addition, neither a transient reduction nor recovery in FNS in G5G8^LKO-A^ mice was observed. Therefore, we limited our subsequent analysis to Control and G5G8^LKO^ mice.

Next we reasoned that if biliary cholesterol is disrupted and less cholesterol arrives in the small intestine via the common bile duct, TICE must close this cholesterol excretion gap somewhere along the length of the intestinal tract. Food consumption and biliary lipid output are entrained to the light/dark-fasting/feeding cycle. Ileal Fgf15 levels are greatest 6 hours after lights off during the post prandial phase indicating active biliary cholesterol secretion and delivery of hepatic bile to the small intestine at this time (31). We euthanized a cohort of mice at this time to assess cholesterol content along the length of the GI tract across genotypes. The intestinal segments and their contents were excised, weighed, and immediately placed in Folch reagent for lipid extraction. While cholesterol content declined along the length of the intestinal tract, it did not differ in any segment between genotypes (Figure 2A). These data indicate that the “sterol gap” within the small intestine due to reduced biliary contribution is closed in the proximal segment of the small intestine.

**Figure 2.**
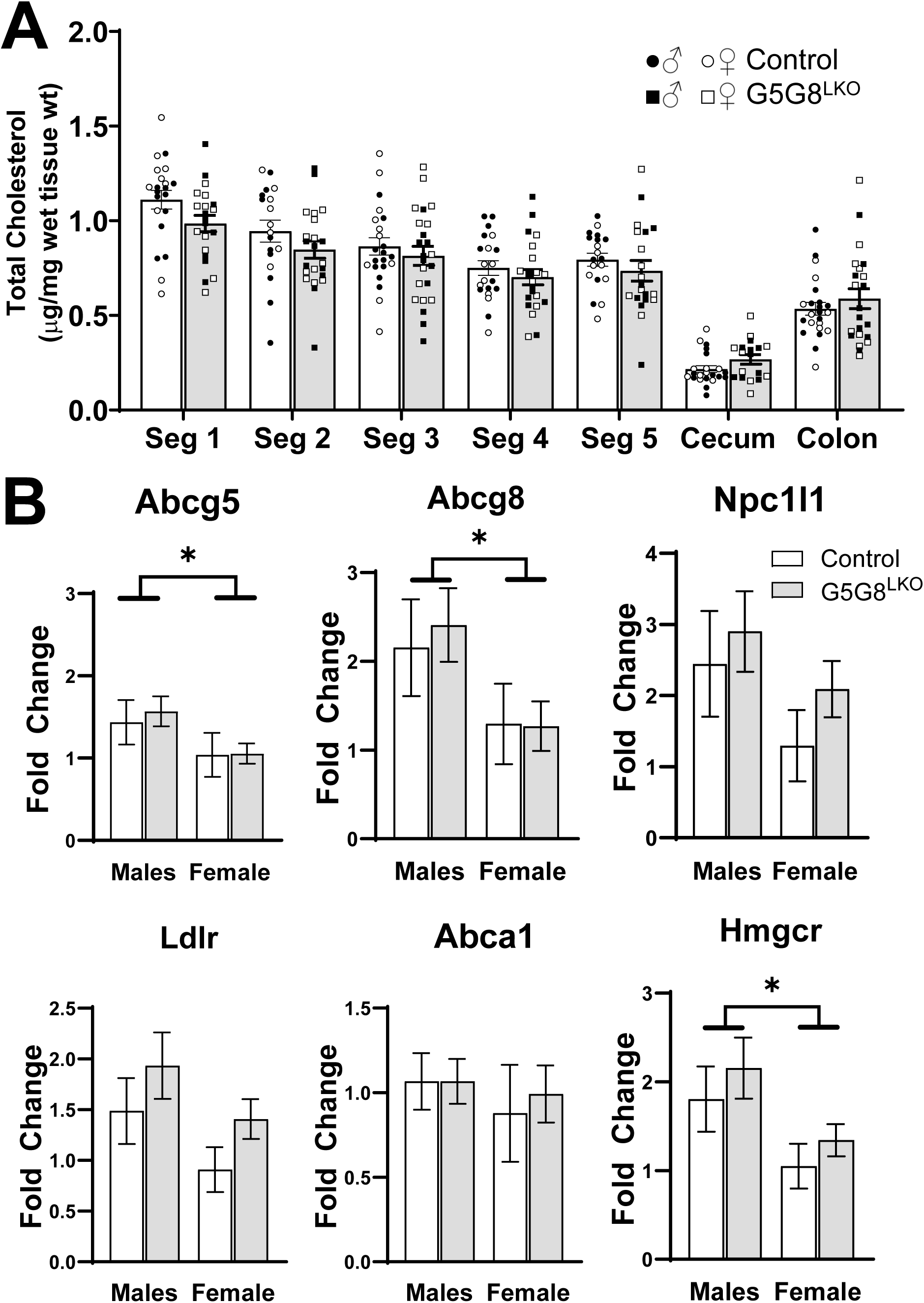
Tissue and lumenal cholesterol content along the gastrointestinal tract in Control and G5G8^LKO^ mice. A) Mice were maintained on standard diet for 12 weeks of age and euthanized in the post prandial phase six hours following “lights-off”. The small intestine (5 equidistant segments), cecum, and colon were dissected and weighed. Sterols were immediately extracted from whole tissue and lumenal contents and quantified by GC/MS. Areas under the curve for cholesterol were normalized to the internal standard and cholesterol content expressed as µg/mg wet tissue weight. Data are mean ± SEM in both males (filled symbols) and females (open symbols) and were analyzed by two-way ANOVA using segment and genotype as factors with Sidak’s multiple comparisons tests for effect of genotype within each segment (n=20-23). B) The relative abundance of mRNAs encoding sterol transporting enzymes in the proximal small intestine as determined by rtPCR. Data are mean ± SEM and were analyzed by two-way ANOVA using genotype and sex as factors (n=8-12). Bars terminating in horizontal lines indicate main effect of sex irrespective of genotype. *P<0.05

We next evaluated the mRNA expression of sterol transporting enzymes in the first segment of the small intestine in both male and female mice (Figure 2B). While differences were observed between sexes, the abundance of intestinal Abcg5 and Abcg8 mRNA was unaffected by the loss of G5G8 in the liver. Similarly, mRNA expression of the apical sterol uptake transporter, Npc1l1, was unaffected. The mRNA levels of mediators of basolateral uptake, Ldlr, and efflux, Abc1a, were unaltered. Transcripts encoding Hmgcr were not affected, suggesting no major shift in sterol synthesis. While there is an apparent trend towards increased Npc1l1, Ldlr, and Hmgcr consistent with a reduction in cholesterol in the proximal small intestine, none of these differences reached statistical significance within or irrespective of sex (i.e. Ldlr_Genotype P=0.15),

The loss of hepatic G5G8 is expected to promote the accumulation of cholesterol in the liver, reduce synthesis and increase its metabolism to primary bile acids. The expression of both Hmgcr and Hmgcs are elevated in female mice relative to males and are reduced in G5G8^LKO^ mice irrespective of sex (Figure 3). However, mRNA for Ldlr was unaffected. We previously published that the loss of G5G8 promoted de novo lipogenesis in mice fed a high fat diet (32). Srebf1 and its target genes Fasn and Acc1 were not affected by the loss of hepatic G5G8 in mice maintained on standard diet. The expression of the rate-limiting enzymes in bile acid synthesis, Cyp7a1 and Cpy8b1, differed by sex but were not affected by the absence of G5G8.

**Figure 3.**
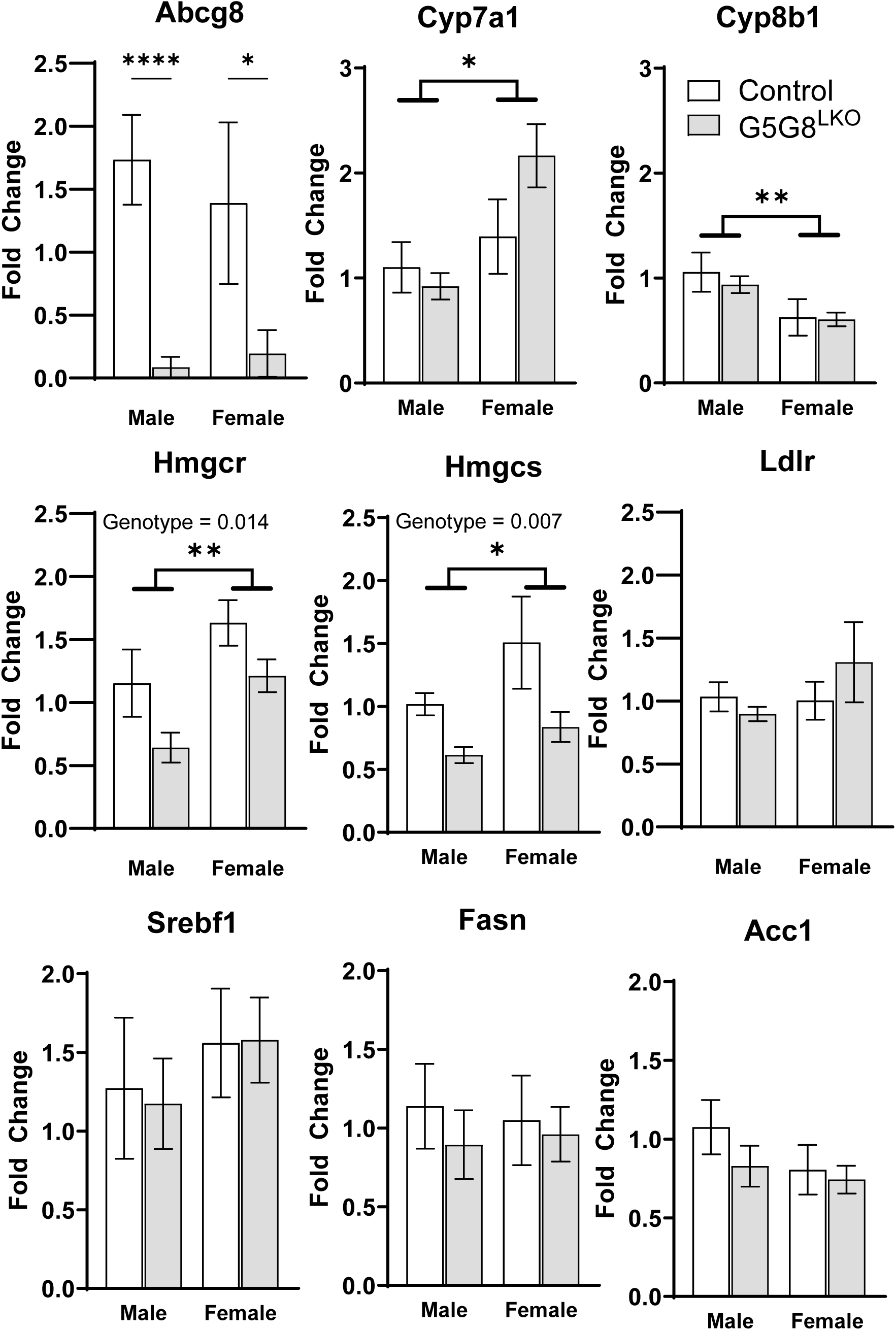
Expression of selected enzymes in lipid metabolism and synthesis in the liver of Control and G5G8^LKO^ mice. Total RNA was isolated from the liver of Control and G5G8^LKO^ mice (Figure 1) and the relative abundance of mRNA of selected enzymes quantified by rtPCR. Data are mean ± SEM and analyzed by two-way ANOVA using genotype and sex as factors as with Sidak’s multiple comparisons tests for effect of genotype within each sext (n=8-12). Bars terminating in horizontal lines indicate main effect of sex irrespective of genotype. P-values for effect of genotype irrespective of sex are indicated by text. *P<0.05 **P<0.01 ***P<0.001, ****P<0.0001

To determine if there was a compensatory upregulation in intestinal cholesterol secretion associated with the loss of hepatic G5G8 and biliary cholesterol secretion, we employed our previously published method for simultaneous determination of biliary and intestinal cholesterol secretion (15, 24). The rate of biliary cholesterol secretion was reduced in both male and female mice G5G8^LKO^ mice relative to their littermate controls (Figure 4A, B). However, we observed no increase in intestinal cholesterol secretion in either sex. We and others who have examined rates of intestinal cholesterol secretion under a variety of conditions have done so using micelles prepared in the absence of cholesterol (Cholesterol Free). A shortcoming of this approach is that bile is not cholesterol free, even in the absence of hepatic G5G8. Therefore we prepared mixed micelles that contained cholesterol within the physiological range (Cholesterol Enriched, Figure 4 C,D). Again, we observed a substantial reduction in biliary cholesterol secretion in the absence of any change in secretion from the intestine. The rate of cholesterol perfusion is depicted with a dashed-line and reflects the theoretical rate of cholesterol appearance in the perfusate after passing through the proximal small intestine had there been no net exchange of cholesterol (Figure 4C, D). Under the conditions of the present study, we observed a net loss of cholesterol from the perfusate indicating uptake of cholesterol as the perfusate passed through the the proximal small intestine. None-the-less, genotypic differences were not observed.

**Figure 4.**
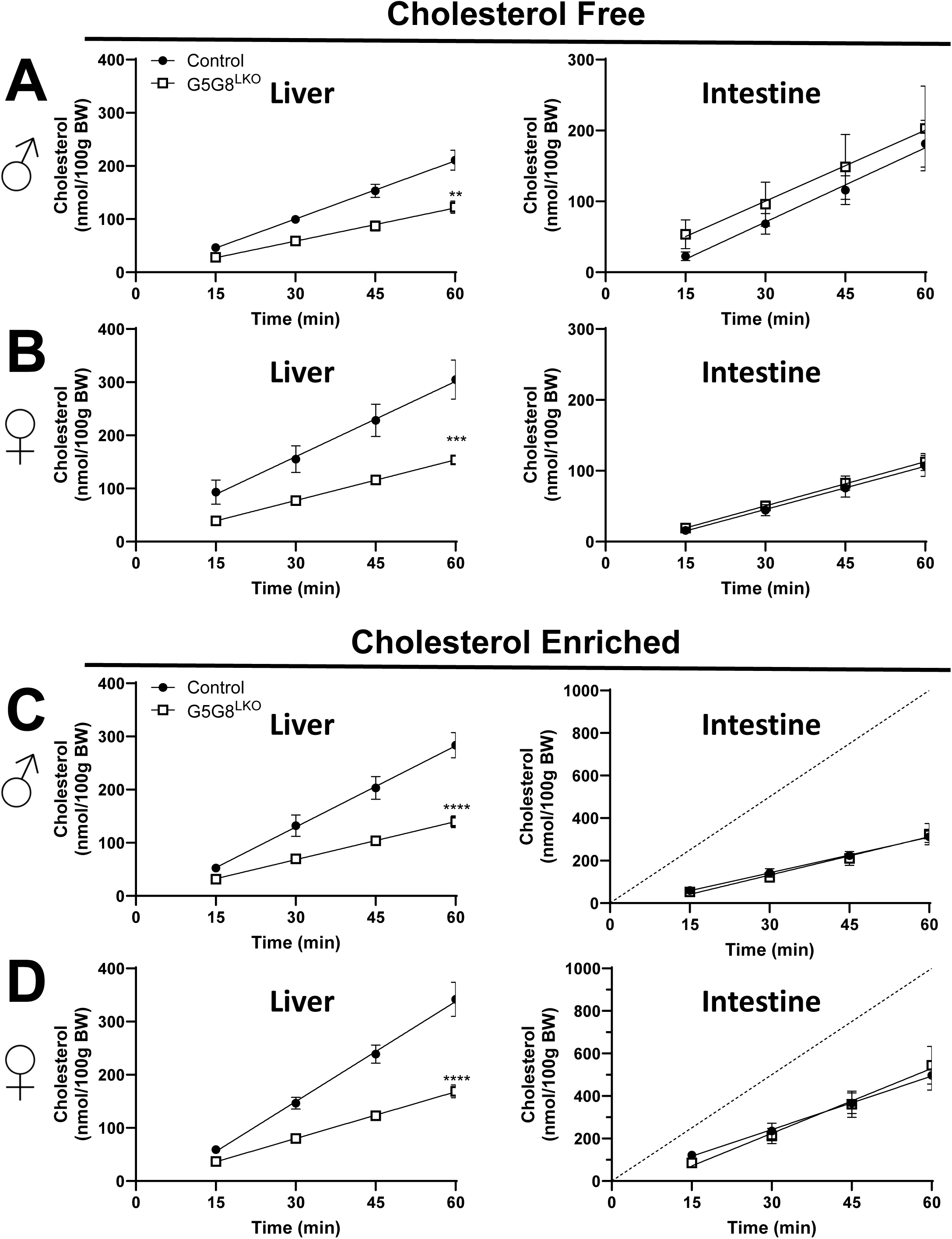
Biliary and intestinal cholesterol secretion in Control and G5G8^LKO^ mice. Mice were bred and housed as in Figure 1. The common bile duct was ligated, the gall bladder cannulated, and bile diverted into collection tubes in both male (A,C) and female (B,D) mice. Bile flow was maintained by continuous tail-vein infusion of TC (100 nmol/min). The proximal small intestine (10 cm) was perfused with Krebs-Henseleit buffer containing mixed micelles (10 mM TC, 2 mM PC) prepared in the absence (A-B, Cholesterol Free) or presence of cholesterol (C-D, Cholesterol Enriched, 0.32 mM). Bile was collected in 15-minute intervals and flow determined gravimetrically. Cholesterol content of bile and perfusates were determined by colorimetric assays and the cumulative rates of cholesterol secretion from the liver (left) and intestine (right) were calculated. The dashed line reflects the rate of cholesterol perfusion (C, D). Data are mean ± SEM and were analyzed by linear regression. Statistical differences in slopes are indicated (n=8-10). **P<0.01 ***P<0.001, ****P<0.0001

Prior studies of G5G8-independent cholesterol secretion were conducted in mice lacking G5G8 in both liver and intestine. Therefore, we also performed these experiments in whole-body G5G8-deficient mice and their wild-type littermates (Figure 5). Our whole-body G5G8-deficient strain is maintained on a PS-free diet in order to prevent confounding effects of biologically active phytosterols present in standard rodent cereal-based diets that vary within and between suppliers as well as to improve breeding efficiency (32, 33). Reductions in biliary cholesterol secretion were observed in G5G8^-/-^ mice in both sexes regardless of intestinal perfusate. While the regression lines for intestinal cholesterol secretion are not superimposable between genotypes, none of these apparent differences reached statistical significance.

**Figure 5.**
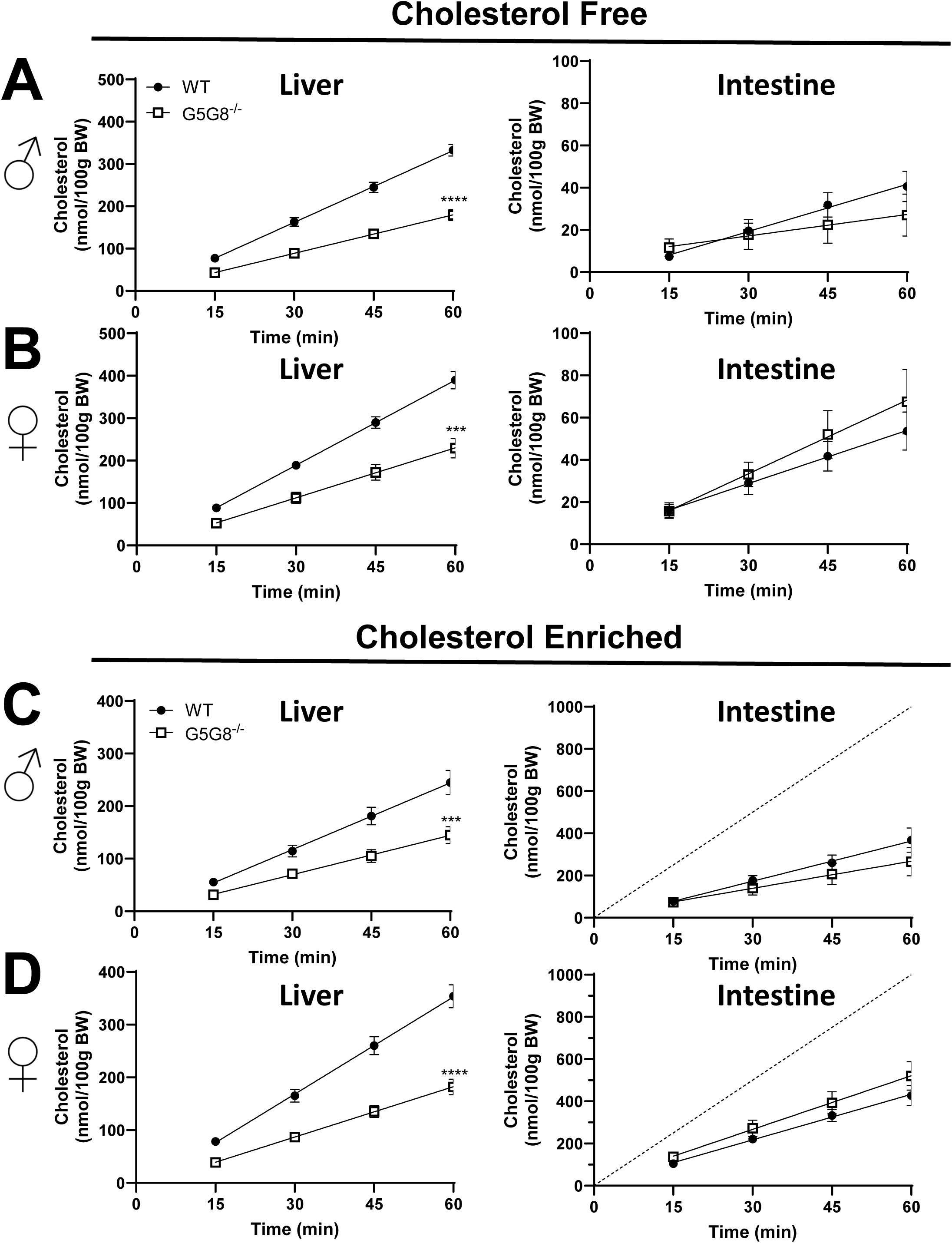
Biliary and intestinal cholesterol secretion in wild-type and whole-body ABCG5 ABCG8 deficient mice. Mice maintained on plant-sterol free diets and analyzed at 8-10 weeks of age. The common bile duct was ligated, the gall bladder cannulated, and bile diverted into collection tubes in both male (A, C) and female (B, D) mice. Bile flow was maintained by continuous tail-vein infusion of TC (100 nmol/min). The proximal small intestine (10 cm) was perfused with Krebs-Henseleit buffer containing mixed micelles (10 mM TC, 2 mM PC) prepared in the absence (A-B, Cholesterol Free) or presence of cholesterol (C-D, Cholesterol Enriched, 0.32 mM). Bile was collected in 15-minute intervals and flow determined gravimetrically. Cholesterol content of bile and perfusates were determined by colorimetric assay and the cumulative rates of cholesterol secretion from the liver (left) and intestine (right) were calculated. The dashed line reflects the rate of cholesterol perfusion (C, D). Data are mean ± SEM and were analyzed by linear regression. Statistical differences in slopes are indicated (n=8-10). ***P<0.001, ****P<0.0001

Our data fail to support an adaptive response in the proximal small intestine that neither elevates the inherent rate of intestinal cholesterol secretion nor reduces the rate of cholesterol absorption to compensate for reductions in biliary cholesterol secretion in order to maintain sterol balance. We next asked if TICE could maintain sterol balance in the absence of hepatic G5G8 in the face of a dietary cholesterol challenge. We first placed the mice on the PS-Free diet for a period of two weeks and measured fecal neutral sterols (Figure 6A). As on standard rodent diet, the absence of hepatic G5G8 had no impact on fecal neutral sterol excretion. Following two weeks on the same diet supplemented with cholesterol (0.2% w/w), control mice increased fecal neutral sterol output by 8-fold. G5G8^LKO^ mice appear to partially compensate for the increased dietary cholesterol (3.37 ± 0.48 vs. 8.40 ± 2.11), but the difference between FNS on the PS-Free diet compared to the cholesterol supplemented diet failed to reach statistical significance. Plasma cholesterol was not affected by the loss of hepatic G5G8 on either diet, but was elevated in the liver of G5G8^LKO^ mice compared to controls irrespective of sex and statistically significant in males following the dietary cholesterol challenge (Figure 6B,C).

**Figure 6.**
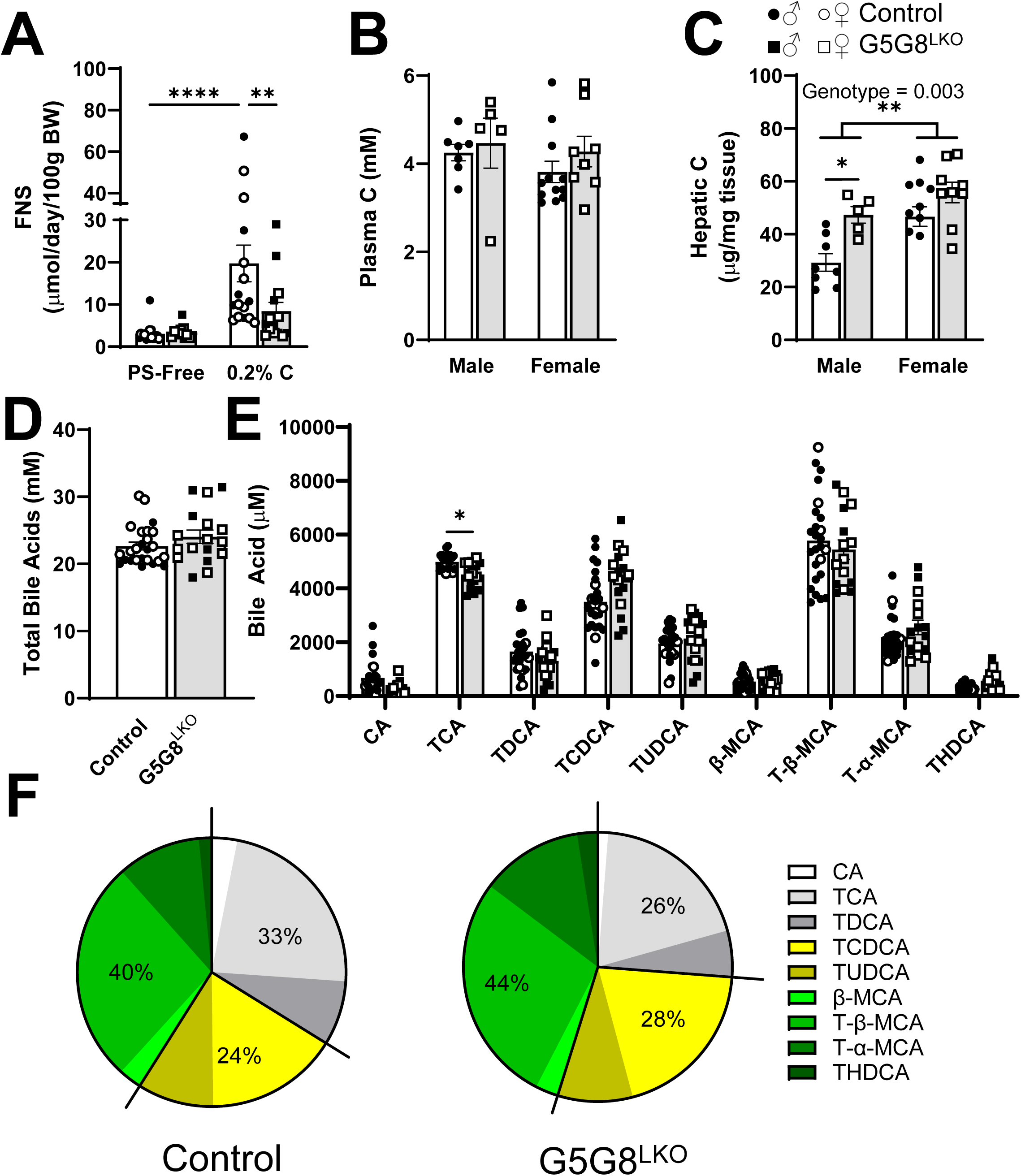
Fecal neutral sterols and bile acid composition of basal bile collected from Control and G5G8^LKO^ mice in response to alterations in dietary cholesterol. Mice were bred and housed as in Figure 1. At 10 weeks of age, mice were switch to the PS-Free diet for a period of two weeks. Mice were then fed the PS-Free diet supplemented with cholesterol (0.2% wt/wt) for a period of two weeks. Feces were collected over the final 3-days of each diet. Basal bile (30 min) was collected and tissues dissected and snap frozen until analysis. A) Fecal neutral sterols in mice maintain on control and cholesterol-supplemented diet. Total cholesterol was also determined in terminal plasma (B) and liver (C). Bile acids were analyzed by LC-MS/MS. D) All detected bile acids were summed to calculate the concentration of total bile acids. E) Concentrations of individual major bile acids (>1% total) and their metabolites. F) The contribution of the major bile acid classes (cholates, chenodeoxycholates, and muricholates) to the bile acid pool in basal bile. Data are mean ± SEM (n=6-12) for measures in both male (filled symbols) and female (open symbols) mice and were analyzed by by two-way ANOVA using diet and genotype (A), sex and genotype (B-C), or species and genotype (E) as factors with Sidak’s multiple comparisons tests for effects of genotype. Bars terminating in horizontal lines indicate main effect of sex irrespective of genotype. P-values for effect of genotype irrespective of sex are indicated by text. *P<0.05, **P<0.01, ***P<0.001, ****P<0.0001

We next evaluated the bile acid pool in basal bile. Total bile acids were unaffected by hepatic G5G8 deficiency (Figure 6D). The composition of the bile acid pool was modestly affected by a small reduction in TC (Figure 6E). Total cholates were slightly reduced (33% vs. 26%) in G5G8LKO mice and offset by small increases in both chenodeoxycholates and muricholates (Figure 6F).

Finally, we analyzed FNS output in wild-type and G5G8^-/-^ mice at baseline (PS-Free diet), following two weeks on the cholesterol-supplemented diet, and two weeks on standard rodent diet. The absence of G5G8 had no effect on FNS output on PS-free diet (Figure 7A). WT mice increased FNS by 33-fold (1.12 ± 0.05 vs. 33.78 ± 3.37, mean ± SEM) following cholesterol feeding while G5G8^-/-^ mice failed to do so (0.96 ± 0.33 vs 3.2 ± 0.179). Following two-weeks on standard diet, genotypic differences in FNS output were no longer observed.

**Figure 7.**
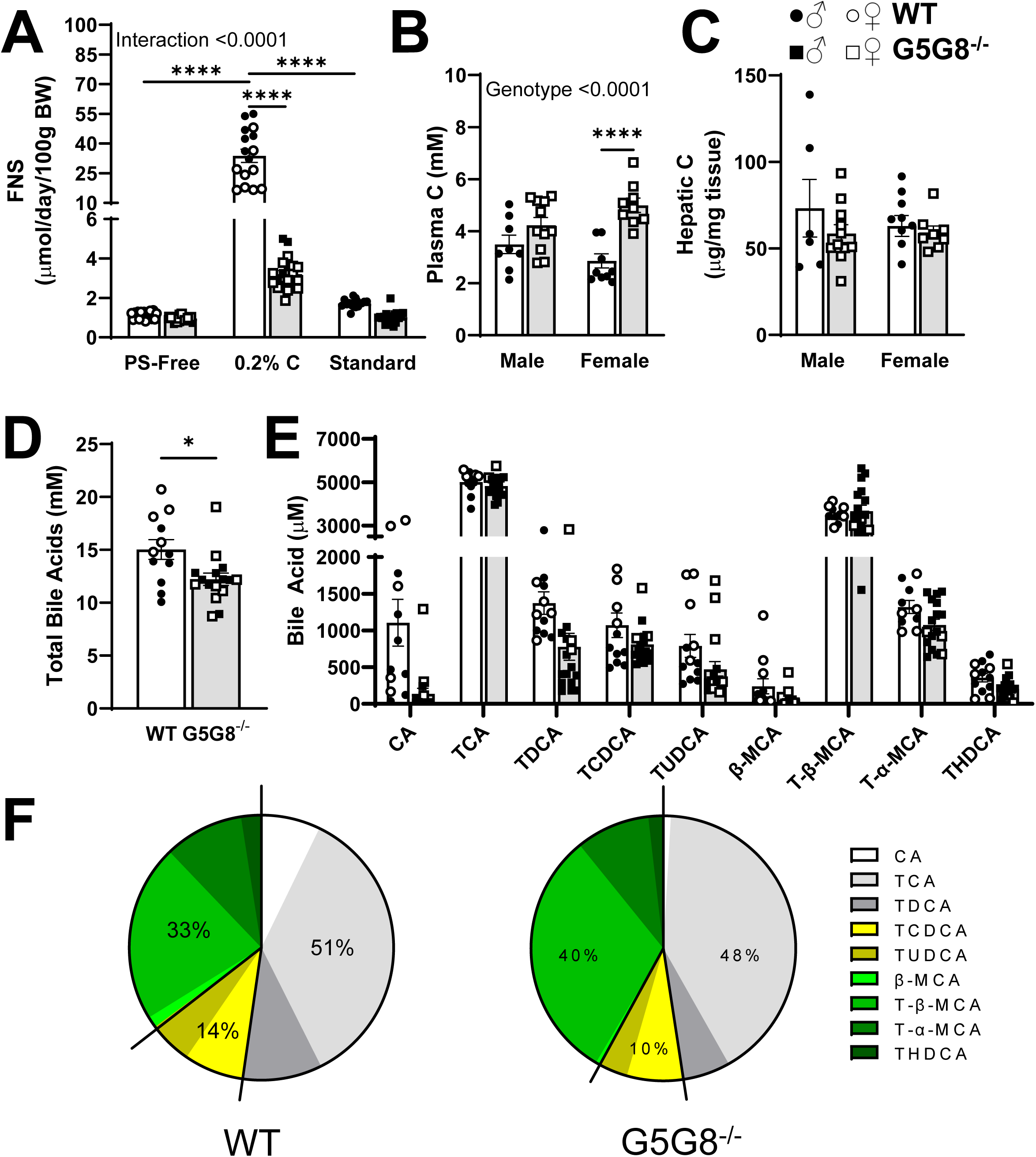
Fecal neutral sterols and bile acid composition of basal bile collected from wild-type and G5G8^-/-^ mice maintained on multiple diets. Mice were bred and maintained on PS-Free diet up to 10 weeks of age. Baseline feces were collected for 3 days before mice were switched to PS-Free diet supplemented with cholesterol (0.2% wt/wt) for a period of two weeks. Mice were then switched to standard diet for a period of 2 weeks. Feces were collected over the final 3-days of each diet. Basal bile (30 min) was collected, tissues dissected, and snap frozen until analysis. A) Fecal neutral sterols in mice maintain on indicated diet were determined by GC-MS. Total cholesterol was also determined in terminal plasma (B) and liver (C). Bile acids were analyzed by LC-MS/MS. D) All detected bile acids were summed to calculate the concentration of total bile acids. E) Concentrations of individual major bile acids (>1% total) and their metabolites. F) The contribution of the major bile acid classes (cholates, chenodeoxycholates, and muricholates) to the bile acid pool in basal bile. Data are mean ± SEM (n=6-12) for measures in both male (filled symbols) and female (open symbols) mice and were analyzed by two-way ANOVA using diet and genotype (A), sex and genotype (B-C), or species and genotype (E) as factors with Sidak’s multiple comparisons tests for effects of genotype. P-values for effect of genotype irrespective of sex are indicated by text. *P<0.05, ****P<0.0001

Terminal plasma cholesterol was elevated in G5G8^-/-^ mice compared to WT regardless of sex and statistically significant in females despite two weeks on standard rodent diet. Hepatic cholesterol was unaffected by genotype following two weeks on standard diet. It should be noted that these terminal measures were conducted with enzymatic colorimetric assays that fail to distinguish cholesterol from phytosterols that likely accumulate in G5G8^-/-^ mice fed a standard diet. The total bile acid pool in basal bile was slightly lower in G5G8^-/-^ mice, but bile acid composition was largely unaffected (Figure 7D-F). There was a modest increase in total muricholates in G5G8^-/-^ mice at the expense of cholates and chenodeoxycholates.

## Discussion

The present series of experiments is consistent with the prior literature demonstrating that a reduction in biliary cholesterol secretion has no effect on cholesterol balance in mice maintained on standard diets. However, our results do not support a compensatory upregulation of TICE as an adaptive response to compromised biliary cholesterol secretion. Intestinal cholesterol secretion rates were unaffected by the loss of G5G8 in liver or in both liver and intestine. Further, we observed no reduction in intestinal cholesterol content along the length of the small intestine in mice lacking hepatic G5G8. van de Peppel et al. demonstrated bidirectional flux of cholesterol in which a substantial portion of cholesterol secreted by the intestine is efficiently reabsorbed, indicating competing absorptive/reabsorptive and secretory processes across the intestinal epithelium (34). We conclude that the arrival of cholesterol-poor bile in the proximal small intestine favors the loss of cholesterol from intestinal enterocytes and closes the gap in intestinal sterol content due to the reduction in biliary cholesterol secretion.

The constant rate of intestinal cholesterol secretion was observed between genotypes regardless of the presence of cholesterol in mixed micelles in the intestinal perfusate. It should be noted that in the experiment using cholesterol-enriched perfusates, less cholesterol exited the first 10 cm of the small bowel than entered. Thus, a fraction of cholesterol was taken-up by the intestinal epithelium indicating net absorption. Importantly, the rate of apparent absorption did not differ between genotypes in this acute setting. These results are consistent with our gene expression data. Transcriptionally, we did not observe an increase in the expression of the LDLR or intestinal G5G8 that would be expected to facilitate the transcellular transport of cholesterol across the enterocyte. Similarly, we did not see a reduction in ABCA1 or NPC1L1 that would be expected to reduce the absorptive pathway to favor TICE.

In our time course study of acute liver G5G8 deficiency, we do not know how rapidly genetic inactivation of *Abcg5 Abcg8* resulted in a loss of G5G8 protein and biliary cholesterol secretion. Our first measure of FNS excretion was 3-days following AAV delivery. We observed no difference in FNS at any point up to 28 days. Biliary cholesterol secretion was clearly reduced in bile collected at termination of the experiment. Thus, the adaptation to hepatic G5G8 deficiency is seamless with no apparent period of reduced cholesterol elimination followed by recovery. We also observed no differences in total fecal bile acids or the concentration of bile acids in basal bile. While differences in bile acid composition in both liver-specific and whole-body knockout mice that favor less hydrophobic bile, it is unclear if this difference would be sufficient to impact TICE. In prior perfusion studies of TICE, hydrophilic and hydrophobic bile acids had similar effects on the rates of TICE whereas phospholipid content in the perfusate had a more substantial effect (12). The composition of our model bile did not examine other bile acids which may have some effect on apparent rates of cholesterol secretion.

To this point it has been assumed that TICE can compensate for disruptions in biliary cholesterol secretion. Indeed, we observe this in mice maintained on standard diet and a purified diet containing trace amounts of cholesterol. However, the presence of cholesterol at 0.2% in the diet overwhelmed TICE or other adaptive responses to maintain sterol output and prevent accumulation in plasma and tissues. While TICE is both measurable and can be stimulated pharmacologically, it is incapable of providing a defense against dietary cholesterol in the absence of G5G8. This may or may not be true in humans. Hypercholesterolemia is not always observed in the clinical presentation of *STSL*. This may simply reflect the amount of cholesterol in the diets of *STSL* patients. However, it may be the case that TICE offers some protection from hypercholesterolemia in sitosterolemics provided their diet is sufficiently low in animal fat and doesn’t overwhelm this mechanism.

Stimulating TICE is an attractive target for accelerating RCT and reducing the risk of atherosclerotic cardiovascular disease and in patients with established plaque burden. TICE bypasses the hepatobiliary pathway and would not be expected to increase the risk of gallbladder disease. Given the prominent role of G5G8 in biliary cholesterol secretion, it is not surprising that variants in *ABCG5* and *ABCG8* have also been linked to cholesterol gallstones. While TICE can be stimulated by pharmacological agents, this has not been demonstrated in the absence of G5G8. Genetic and pharmacological approaches for intestinal-specific stimulation of the LXR pathway have been shown to accelerate RCT and reduce atherosclerosis in mouse models. Thus, the stimulation of G5G8-dependent TICE offers a viable path forward to accelerate cholesterol excretion in the absence of increased gallstone risk.

## Data Availability Statement

All data will be retained and made available upon requestion in accordance with institutional policies at the University of Kentucky.

## Acknowledgements

The authors appreciate the assistance of the staff of the Division of Laboratory Animal Resources at the University of Kentucky in the maintenance and care of our colony.

---

**Table S1.**
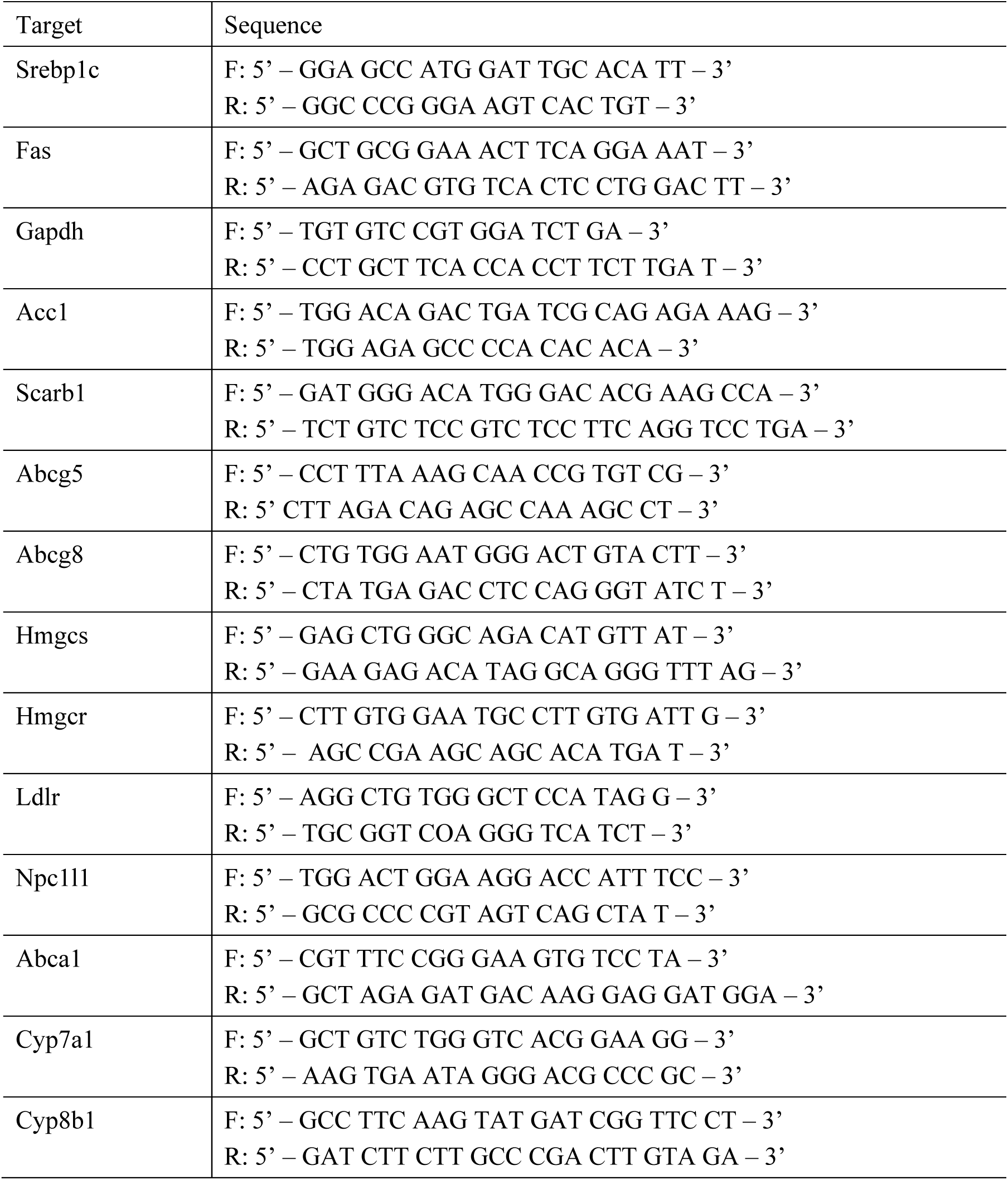
Primer Sequences for rtPCR analysis of transcript abundance.

